# Mutation divergence over space in tumour expansion

**DOI:** 10.1101/2022.12.21.521509

**Authors:** Haiyang Li, Fengyu Tu, Lijuan Deng, Zixuan Yang, Yuqing Han, Xing Fu, Long Wang, Di Gu, Benjamin Werner, Weini Huang

## Abstract

Mutation accumulation in tumour evolution is one major cause of intra-tumour heterogeneity (ITH), which often leads to drug resistance during treatment. Previous studies with multi-region sequencing have shown that mutation divergence among samples within the patient is common, and the importance of spatial sampling to obtain a complete picture in tumour measurements. However, quantitative comparisons of the relationship between mutation heterogeneity and tumour expansion modes, sampling distances as well as the sampling methods are still few. Here, we investigate how mutations diverge over space by varying the sampling distance and tumour expansion modes using individual based simulations. We measure ITH by the Jaccard index between samples and quantify how ITH increases with sampling distance, the pattern of which holds in various sampling methods and sizes. We also compare the inferred mutation rates based on the distributions of Variant Allele Frequencies (VAF) under different tumour expansion modes and sampling sizes. In exponentially fast expanding tumours, a mutation rate can always be inferred in any sampling size. However, the accuracy compared to the true value decreases when the sampling size decreases, where small sampling sizes result in a high estimate of the mutation rate. In addition, such an inference becomes unreliable when the tumour expansion is slower such as in surface growth.

## Introduction

The accumulation of somatic mutation drives the tumour growth and development (Abascal *et al*., 2021). The higher level of the heterogeneity of tumour tissue, the higher the difficulty of treatment and the higher the recurrence rate in clinical practice (Sun & Yu, 2015; Turajlic *et al*., 2019). While it is a complicated process driven by both spatial and temporal dynamics (Akdemir *et al*., 2020), tracking and analysing the distribution of accumulated mutations is of great significance to understanding intra-tumour heterogeneity (ITH) (Martincorena *et al*., 2018; Turner & Reis-Filho, 2012; Greaves & M, 2015). While temporal samples in clinical are often infeasible or not available, more and more spatial studies of ITH have been carried out by analysing sequencing data sampled at different locations (de Bruin *et al*., 2014; Joshi *et al*., 2019; Bashashati *et al*., 2013; Blokzijl *et al*., 2016; Ryser *et al*., 2018). Combining with various computational approaches, evolutionary parameters such as selection strength and the time of tumorigenesis were inferred.

Sottoriva *et al*. (2015) introduced a ‘Big Bang’ model and showed that early mutations after tumorigenesis can be neutral without bringing additional fitness advantages to the initial tumour cells across many cancer types. The distribution of those early mutations follows a power law decay along the increase of mutation frequency (Williams *et al*., 2016). Waclaw *et al*. (2015) described another growth model of exponentially expanding tumours and showed that the ITH level and selective advantage have a negative correlation, and high heterogeneity leads to the rapid onset of resistance to chemotherapy. In addition, even a small amount of localized cellular dispersal can cause a faster growth of tumours formed of conglomerates of balls with more similar genetic alterations, compared to spherical tumours without cell dispersal. Similarly, Ryser *et al*. (2018) investigated the role of early tumour cell migration in shaping the patterns of private mutation in colorectal tumours by simulating exponential growth in the founder gland followed by consecutive gland fission. Their results showed that detecting the same private mutations in opposite sides of the final tumours indicates early cell mixing and movement, and that neutral evolution leads to high local ITH under the single expansion hypothesis of tumorigenesis.

Beside of exponential growth, Lenos *et al*. (2018) and Fu *et al*. (2022) also visualized the heterogeneity distribution and clonal diversity under volume and surface growth. Lenos *et al*. (2018) quantitative analyses of lineage tracing data from primary colon cancer xenograft tissues. They found out that the stem cells driving tumour growth are mostly at tumour edges, and stem cell properties change over time depending on the cell location. Fu *et al*. (2022) showed that the complete variability of clone size is due to spatio-temporal regulation of necrosis and that surface growth had a greater effect on sub-clonal diversity than volume growth. They also revealed that clonal diversity decreased sharply with the growth time in the surface model. Gallaher *et al*. (2018) applied the off-lattice agent-based model to systematically study and predict the spatial dynamic evolution of heterogeneous tumours in different treatment periods. Their results showed that evolution-based strategies by exploiting the cost of resistance can delay treatment failure. Meanwhile, fewer drugs and more vacation-oriented treatment can reduce tumour heterogeneity.

While different growth modes of solid tumours are likely to be related to the strength of spatial constrains on tumour growth and tissue-specific (West *et al*., 2021), we model a continuous inter-cellular push rate (0 ≤ *p* ≤ 1, Chkhaidze *et al*., 2019), i.e. the likelihood cells push neighbour cells outward when there is no direct empty space around during expansion (Radszuweit *et al*., 2009; Waclaw *et al*., 2015; Sottoriva *et al*., 2017; Ryser *et al*., 2018; Chkhaidze *et al*., 2019). Different values of push rate refer to different modes of tumour growth including the surface (*p* = 0) and exponential growth (*p* = 1) as two boundary examples. We are interested in how the spatial constrain and sampling distance will impact on the measured ITH between samples.

Recent studies show that sampling can have an important impact on the measured ITH. Opasic *et al*. (2019) simulated tumour heterogeneity in a lattice model and investigated how sampling will impact on the accuracy of identifying clonal mutations. They showed non-random sampling from a circular spatial pattern might improve the classification of true clonal mutations compared to random sampling. Ling *et al*. (2015) evaluated the spatial distribution of point mutations by sequencing about 300 sampled regions within the circular range of a single tumour. Combining with an exponential growth model, they found out that cells sampled farther away from the sample centre carry more mutations. Moreover, other experimental studies also indicated that cell density and metastatic likelihood differ between tumour centre and margin (Masugi *et al*., 2019; Zhao *et al*., 2021). Here, we take random samples with various sizes multiple times in each simulation and analysis the average pattern of ITH between samples. In comparison, we also take multiple samples from the central 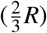 and marginal regions 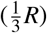 of simulated tumours with radius *R*. We record the accumulation of point mutations, compare mutation frequencies above a detection limit and infer mutation rates in each sample. To compare the ITH between samples, we apply the Jaccard index to quantify the mutation composition difference, which has been used to quantify mutation diversity in much genetic data (Alioto *et al*., 2015; Silva *et al*., 2017; Araf *et al*., 2018; Gendoo *et al*., 2019; Yan *et al*., 2019; Pereira *et al*., 2021). The more mutations shared between two samples, the higher the Jaccard index is and the lower the ITH is.

Our results show that under a given tumour expansion mode (fixed push rate), the ITH increases with the sampling distance, which holds for various sampling sizes and for random, centre and margin sampling. The faster tumour expands (larger push rates), the slower the ITH increases along the sampling distance. Using the recorded mutation information in cells, we construct the distributions of Variant Allele Frequencies and infer the mutation rate for each sample. When push rates are large such as in exponentially expanding tumours, a linear relation is always observed between the number of mutations and accumulative mutation frequencies independent of sampling sizes. Using this linear relation, we estimate the mutation rates and our results show that the sampling size strongly impacts the inferred mutation rate. Smaller sampling sizes lead to an overestimate of inferred mutation rates as well as the larger variance of inferred mutation rates among samples. Increasing the sampling size improves the inference with a more accurate mean compared to the true value of the mutation rate and smaller variance. When the tumour expansion is slower such as in surface growth, such inference becomes impossible, because the linear relationship does not hold. The small the sampling size, the larger deviation the VAF is from a linear relation.

## Methods

### Stochastic simulations of tumour growth in space and mutation accumulation

We simulate tumour growth in a lattice where each cell has eight direct neighbours. Tumours develop from a single cell, which is seeded in the centre of the lattice. When a cell divides into two daughter cells, one cell remains in its original location (*L*_*p*_), and the other daughter cell is located in a randomly selected empty space among its direct neighbours. If there is no free space, with a probability *p* (0 ≤ *p* ≤ 1), a new space can be created by pushing a randomly selected neighbour cell (at location *L*_*d*_) and all the rest cells along the direction of *Lp* and *Ld* outwards for one position until an empty space was reached (Chkhaidze *et al*., 2019). For two extreme values of the push rate, *p* = 0 refers to the surface growth where only cells in the outskirt of the tumour would divide and *p* = 1 refers to an exponential growth where all cells divide in each time step (Fig. 1).

**Figure 1.**
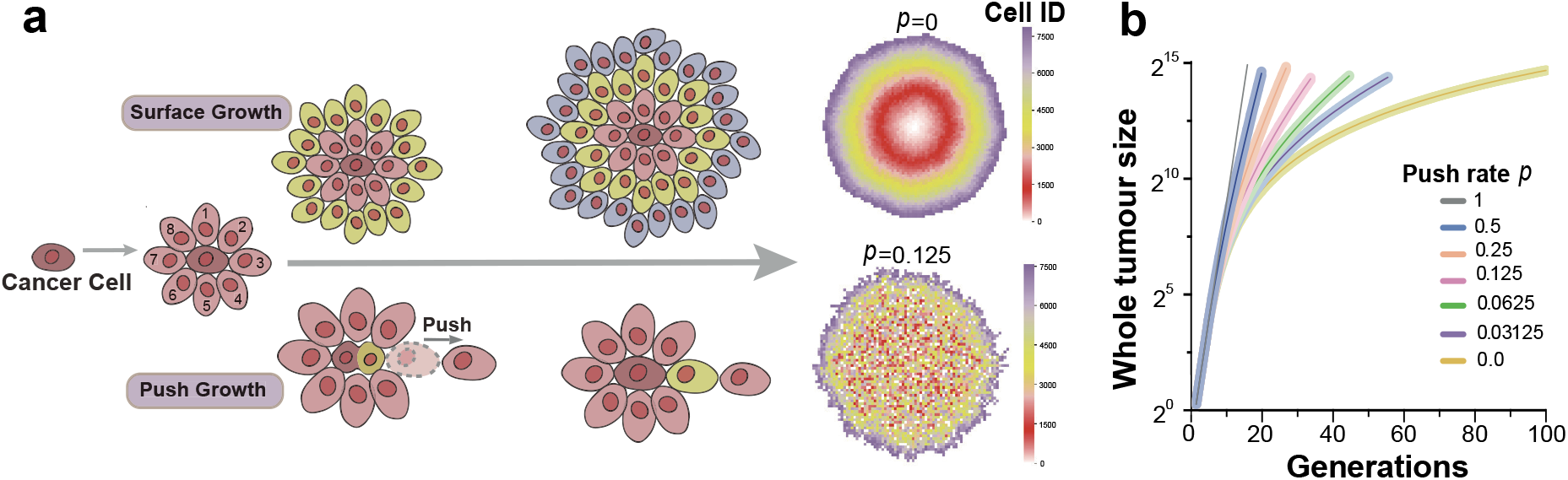
The spatial growth under different push rates. **a**) when push rate *p* = 0, it refers to the surface growth, where only cells on the surface have an empty spot in its direct neighbours and can divide. For other values of the push rate, cells not on the surface can also divide by creating an empty spot among its neighbours through pushing. We allocated a unique ID for each new cell. The larger the cell ID, the later the cell was reproduced. **b**) To reach the same tumour size, surface growth takes more generations of reproductions compared to exponential growth under *p* = 1. While mutation accumulation only happens in cell divisions in our model, for tumours of the same size, the push rate will impact on the mutation burden in tumours and also the spatial distribution of those mutations. The shadowed area is 100 simulations under the corresponding push rate, where their averages are shown as solid lines. (The fig.1b is 100 times simulation, cell number is around 2^15^, *p* = 1 grow to 2^15^ cells only 15 generations. when *p* = 0 divide 100 generations can grow about 27500 cells)

In each cell division, random point mutations can happen in both daughter cells compared to the parent cell. The number of new mutations in each daughter cell follows a Poisson distribution (Clarke, 1946; Bozic *et al*., 2010; Waclaw *et al*., 2015; Williams *et al*., 2016; Williams *et al*., 2018), where *λ*is the mean value representing the average mutations (mutation rate) per cell division. Thus, the probability to have *k* new mutations in one daughter cell is 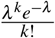. If not specified, we use *λ*= 10, which is conventionally considered as the average number of point mutations per cell division in tumours (Frigola *et al*., 2017; Carlson *et al*., 2018; Williams *et al*., 2018).

### Spatial sampling

To understand how mutations spread over space, we test two different spatial sampling methods, i.e. random sampling and centre-margin sampling. For the random sampling method, we sample 500 locations randomly in each simulated tumour and collect cells in a rectangular area around these locations (Fig. 1a in SI). To investigate the impact of sampling sizes on our diversity measures, we vary each sample size from 100 to 3600 cells (around 0.6% to 21.6% of the whole tumour). Alternatively, we divide simulated tumours into the central region (a circle of 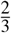 of the radius from the tumour centre) and the marginal region (the rest 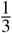 ring structure). We randomly sample 500 rectangular areas with 100 cells in each sample (around 0.6% of the whole tumour) in the margin and 500 samples in the centre region (Fig. 1b, in SI).

### Measurements of intra-tumour heterogeneity between samples

We use a statistical measurement, the Jaccard index, to compare the diversity between samples. Supposing *A* and *B* are the set of mutations in two samples, the similarity of mutations between the two samples is given by

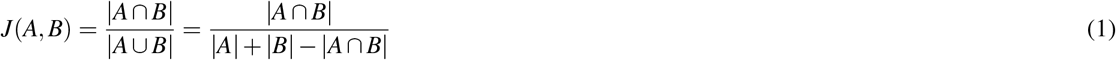

where *A* ∩ *B* is the intersection of mutations in these two samples and *A* ∪ *B* is their union. Thus, by definition the Jaccard index will be between 0 and 1. By using the Jaccard index, we can quantify how similar mutations are accumulated in samples over space. More specifically, we studied how the Jaccard Index is related to the sampling distance, which is measured by the Euclidean distance between the centres of samples.

### Mutation rate inference and Kolmogorov–Smirnov test

Another classical measurement for patterns of mutation accumulation in population genetics is the frequency distribution of mutations among tumour cells in each sample, which is called Variant Allele Frequency (VAF) distribution in cancer research. We are interested in how different growth dynamics, such as push rates as well as spatial sampling impact on the measured VAF distributions. For simplicity, we simulate only one driver events where genetic changes lead to the initiation of a single tumour cell, which seeds for the tumour growth and the accumulation of random neutral mutations during the tumour expansion. In this scenario, the accumulated mutations follow a theoretical expectation of power-law decay, where the number of mutations of a given frequency decrease along with the mutation frequency *f* (Williams *et al*., 2016). The cumulative distribution is a linear relation as 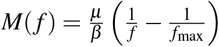, and the slope could determine the effective mutation rate 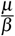 (the true mutation rate scaled by death). We constructed the cumulative VAFs for all samples and compared how the push rates, sampling methods, and sizes would impact on this measurement. We quantify how much the observed cumulative agrees with a linear regression by Kolmogorov-Smirnov (KS) test (Peacock, 1983), which measures the maximum distance of the linear regression with the observed cumulative VAF curve. If the observed cumulative VAFs agree with linear regression, we inferred the mutation rates using the equation above.

## Results

### Spatial mixing increases and variance of mutation frequencies decreases with push rates

When the push rate is low (*p* = 0), cells grow slowly and mainly on the surface. From spatial patterns constructed by the cell ID (Fig.1a), where the cell born later is assigned with a larger ID number, we observed clear circular boundaries among early and later born cells. However, when the push rate increases, these spatial boundaries become loose. Instead, spatial mixing among early and later born cells appears. Similar effects are observed in the spatial pattern of mutations. In Fig.2b, we demonstrate the spatial pattern of four different mutations randomly picked up from four cells born in the second generation. When *p* = 0, clear boundaries among cells carrying those mutations exist. With the increase in push rate, cells are carrying different mutations mix in space.

**Figure 2.**
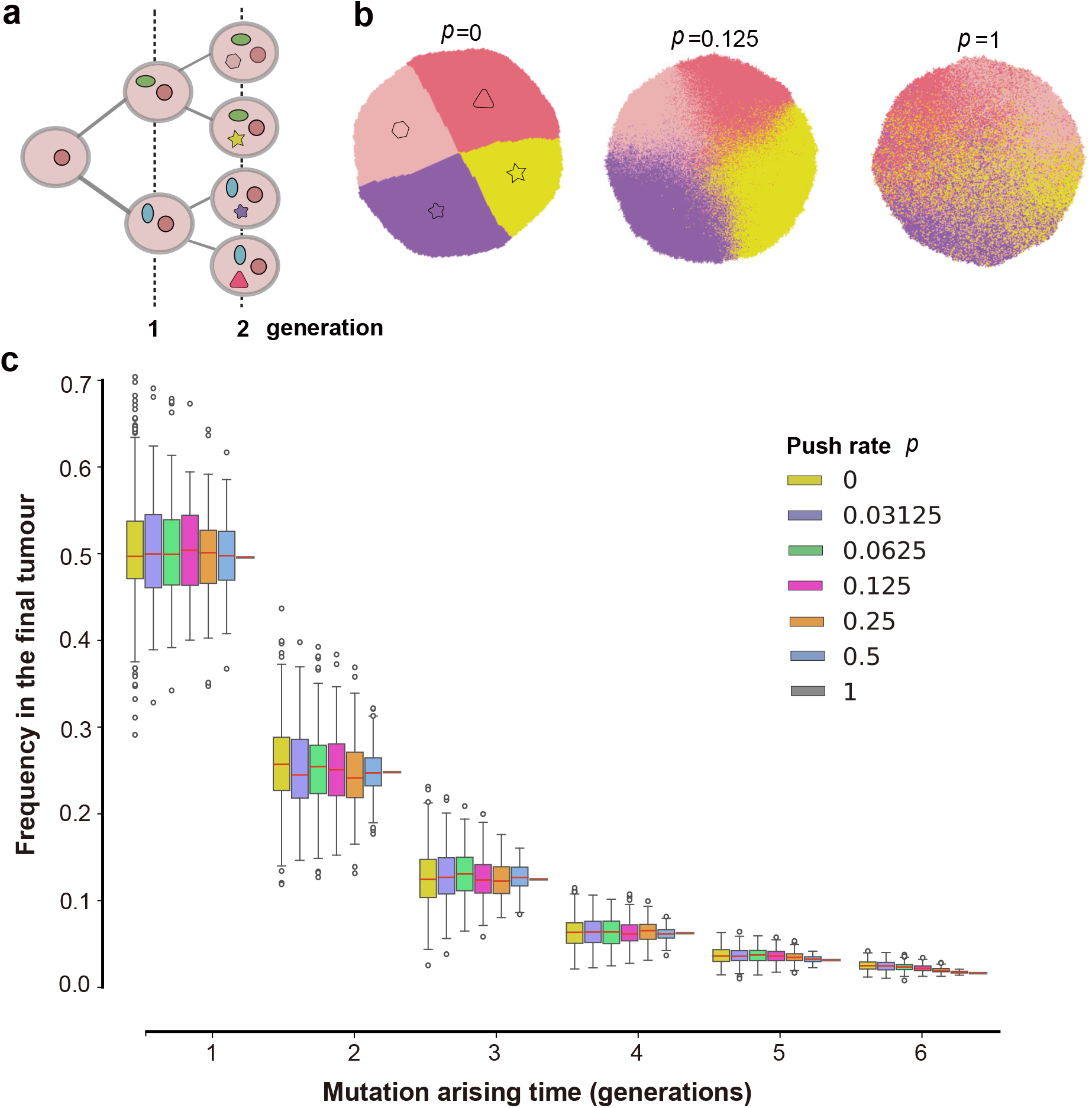
The impact of push rates on distributions of early mutations. **a**) We record the identities of early mutations, including the mutation ID and the time when they were first present in the population. **b**) One example of the spatial distribution of unique mutations from four cells after the second round of cell division. (the distribution of 4 types of mutation in the second generation, cell number is about 2^14^). **c**) Frequencies of all early mutations in growing tumours. These mutations arise at different rounds of early cell divisions (300 times simulation, cell number is about 2^14^, cut last 3 points). While the push rates change, thus the spatial distributions of those mutations differ as in b); their final frequencies are 0.5, 0.25, 0.125, 0.0625 and 0.03125 with the push rate. Here, the red lines in colourful bars are the average mean value of frequencies of all mutations from 300 times simulation realisations, and the bars are the variance. (λ = 10, the final tumour size is 2^14^.)

The earlier a mutation arises during the tumour growth, the higher frequency it reaches in the final tumour Fig.2c. This pattern is consistent under all push rates. Arising at the same tumour generation, the mean value of the frequency these mutations can reach in the final tumour is independent of the push rates. However, the variance increases monotonically when the push rate decreases.

### Growth patterns on the VAF distribution and mutation rate inference

Next, we construct the VAF of all mutations accumulated through tumour growth. Mutations with the frequency of less than 0.01 in the final tumour are discarded, as in reality, it is hard to detect such lower frequency mutations in a standard sequencing depth. Fig. 3a and Fig. 3b show examples of single simulations under two boundaries of push rates, *p* = 0 and *p* = 1. Without mimicking sequencing noise in our simulation, when push rate *p* = 1, the VAF distribution is discrete with mutations at frequencies 0.5, 0.25, 0.125, 0.0625 and so on (Fig. 3 an inner panel). The cumulative VAF distribution is a linear line and perfectly agrees with the theoretical expectation under neutral selection. On the contrary, when only surface growth is allowed under *p* = 0, there is a wide and relative continuous VAF distribution covering all intermediate frequencies and even with few mutations exceeding 0.5 (Fig. 3b inner panel), which is the frequency of most clonal mutations in diploid populations. The cumulative VAF distribution under surface growth strongly deviates from a linear relation, which is quantified by the KS distance. The larger the KS distance, the further away the cumulative VAF distribution deviates from a linear regression.

**Figure 3.**
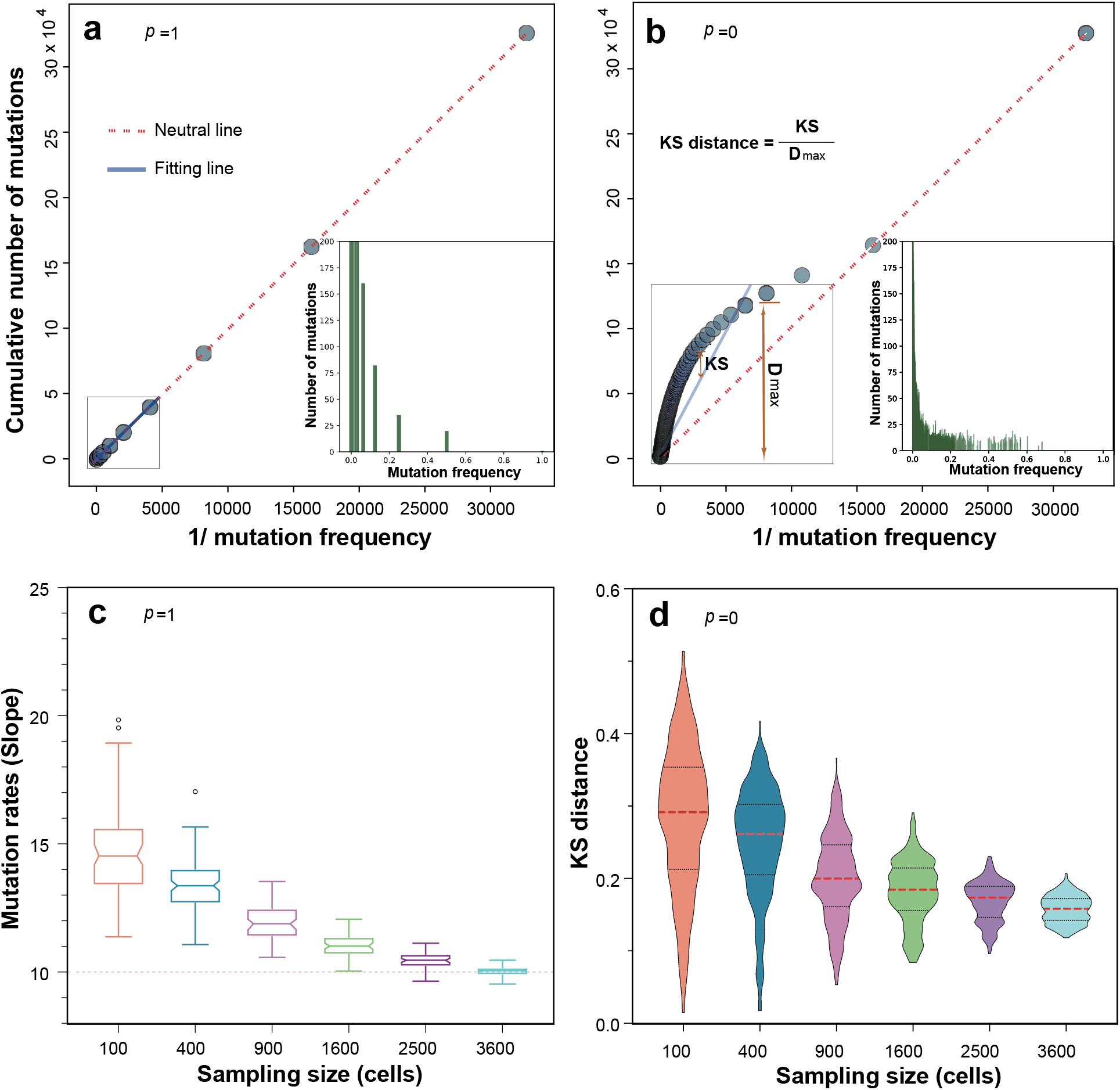
Mutation accumulations under different pushing rates and sampling sizes. **a**) Cumulative distribution of mutations when *p* = 1, where the computational simulations agree with the linear expectation (red solid lines) of neutral mutations accumulated in exponentially expanding tumours. **b**) As the push rate decreases, the tumour growth is closer to the surface growth (*p* = 0). The corresponding cumulative distributions of mutations deviate from the linear expectation, which is measured by the Kolmogorov–Smirnov (KS) distance. **c**) Under exponential growth (*p* = 1), sampling does not change the pattern of linear regression fitting for accumulated mutations. However, the estimated mutation rate increases when the sampling size decreases. **d**) Under surface growth (*p* = 0), the deviation from the linear expectation increases when the sampling size is smaller. (λ = 10, the final tumour size is 2^14^).

We measure the KS distance for different push rates and sampling sizes under random sampling. Note to eliminate the extremely low frequency mutations (less than 0.01), we discard the last few dots in the cumulative VAF distributions for the linear regression (e.g. the last three dots in Fig. 3a and Fig. 3b). We found out the push rates have a strong impact on this measurement, where the KS distance keeps a relatively high level when push rates are small (Fig. 3d and Fig. 4 a-c in the SI). This means that mutation rate inferences based on a linear regression is not reliable under small *p*. The KS distance decreases when *p* becomes larger (Fig. 4 d-e in the SI), and a linear regression is reasonable across all sampling sizes (Fig. 2 d-e in the SI). Thus, we can infer the mutation rates based on the slope of the linear regression under large *p*.

**Figure 4.**
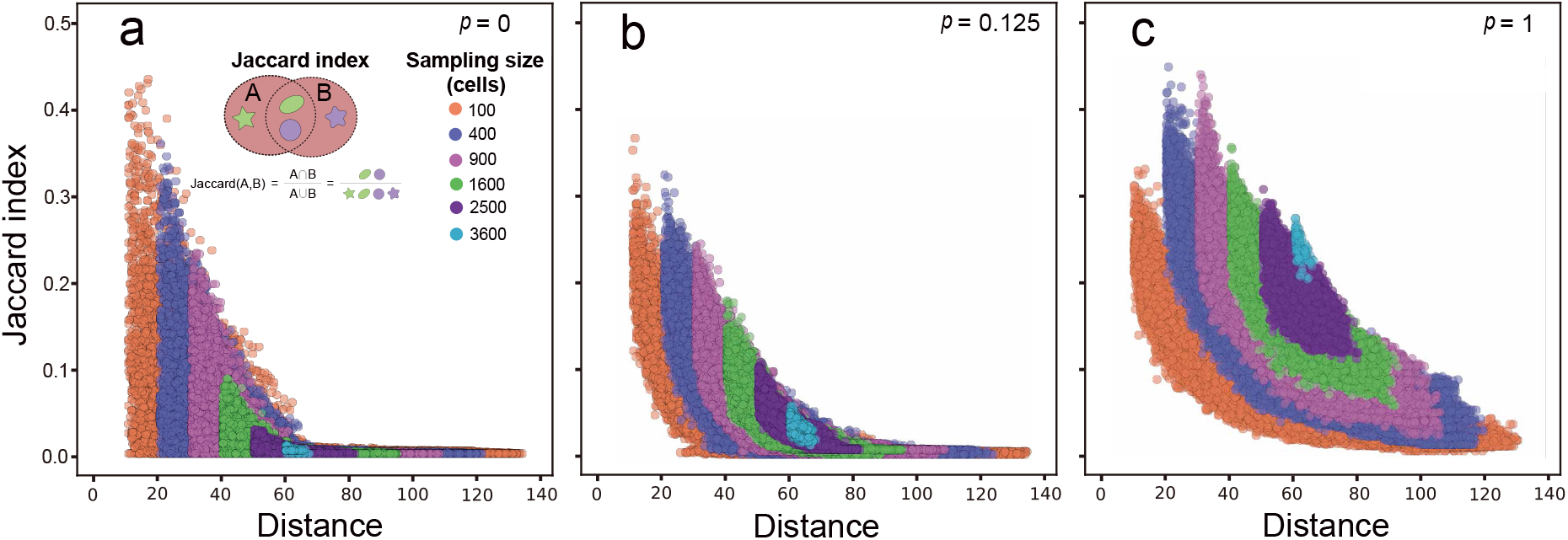
The mutation divergence over space. Pairwise comparison of mutation divergences between samples are measured by Jaccard Index. As the sampling distance increases, fewer mutations are shared between the samples, and the smaller the Jaccard Index is. **a**), *p* = 0. **b**), *p* = 0.125. **c**), *p* = 1. This holds for any push rates and any sampling size. For the surface growth **a**), the Jaccard index decreases faster to 0 when the sampling distance increases compared to exponential growth **c**). This effect is stronger when the sampling size is smaller (e.g. orange dots compared to purple dots). (1-time simulation, sampling 500 points for each sample size, cell number is about 2^14^, cut last 3 points)

Fig.3c shows the fitting results of 100 simulations with 500 random sampling each under *p* = 1. While increasing the sampling size, the inferred mutation rate is closer to the true value with reduced variance. On the contrary, although we also observe a close-to-linear relation under small *p* (Fig. 2 a in the SI), the inferred mutation rate is often an overestimate compared to the true value (Fig.3c, dashed line). In addition, the smaller the sampling size is, the large the variance of the mutation rate is.

### Intra-tumour heterogeneity between samples

In each simulation, we first randomly sample 500 areas with various sampling sizes. We measure the ITH between spatially non-overlapping samples and quantify how this heterogeneity changes with spatial distances. We compare the samples pairwisely to calculate the Jaccard index, which is negatively related to the heterogeneity level. Meanwhile, the spatial distances between samples are defined by the Euclidean distance between the central points of each sample. For various sampling sizes and push rates, the Jaccard index between two random samples decreases, thus the ITH increases, monotonically with the spatial distances (Fig.4, Fig.5 in SI). When the push rate *p* = 0 (surface growth), the Jaccard index drops down to 0 very fast (Fig.4a), where the non-overlapping samples has fewer and fewer shared mutations when the sampling distance increases. When *p* increases, given the same sampling size and distance, the Jaccard index increases. This agrees with the observation of spatial mixing of cells carrying different mutations. When *p* = 1, cell spatial mixing reaches the highest level, and we seldomly observe any Jaccard index as 0 even under the smallest sampling size and largest sampling distance and there are always shared mutations among those samples. While the results in Fig.4 are based on single simulations, 249, 500 Jaccard indexes are calculated after the pairwise combination of 500 samples in a single simulation. Such a large number gives a stable pattern, which is consistent with results over 100 simulations (Fig.6 in SI).

To understand the impact of sampling methods, we divided the simulated tumours into the central region (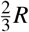 *R* circle) and margin region (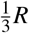 *R* ring width), where *R* is the tumour radius (Fig. 1b in SI). We randomly sampled 500 areas (100 cells, 196 cells, 400 cells) in the margin and centre, respectively (Fig.7 in SI). Then, we compared the relationships of the Jaccard index and spatial distance between samples in the margin and the centre region under the different push rates. The patterns are very similar as those observed in completely random sampling. The Jaccard index decreases with the increase of the sampling distance, and the push rates lead to a higher Jaccard index under the same sampling size. In addition, we see the Jaccard index is slightly higher between samples in the central region compared to the margin region. This is more obvious when push rates are small, where the spatial constrain is stronger and thus less mutations are shared between spatially non-overlapping samples. In summary, the two sampling methods do not alter the qualitative pattern of how ITH increases with the sampling distance and with less spatial constrains (increasing *p*). However, there is a small quantitative difference if we sample in the margin or centre of tumours.

## Discussion

We developed a computational model that tracked the dynamic movement of each cell and variation divergence, which revealed the relationship of spatial heterogeneity distribution with sampling size and tumour expansion modes. We used push rates to model slow and fast growth modes, where small push rates referring to surface growth and large push rates to exponential expansion without spatial constrains. Furthermore, we recorded the mutation accumulation during all growth modes and applied different sampling methods, i.e. completely random sampling and margin-centre sampling, with various sampling sizes.

Under the surface growth (small push rates), the accumulation of mutations is concentrated in a continuous space, and mutations arising in different original cells can form clear boundaries in space. When the push rate increases, the mutations becomes more spatially dispersed, which agrees with the conclusion of Chkhaidze *et al*. (2019) in simulating driver mutations under the action of pushing. We further showed that small push rates introduce high stochasticity in the system. While the final frequency that an early mutation can reach in small push rates has the same mean expectation with large push rates, the variance is much higher. We showed the mutation cumulative VAF distributions agree in a linear relationship with high push rates independent of the sampling size. We can infer mutation rates based on the method proposed in Williams *et al*. (2016). However, a small sampling size will overestimate the mutation rates compared to the true value in our simulations. When the push rate decreases, the mutation cumulative VAF distributions deviate largely from a linear relation, and such a mutation rate inference becomes less meaningful.

While mutation heterogeneity can reveal a tumour’s life history (McGranahan & Swanton, 2017) and the patterns of ITH in space are important to understand and improve tumour treatments, systematical studies to quantify those properties are still rare (Chkhaidze *et al*., 2019; Fu *et al*., 2022). We used the Jaccard Index to quantify the mutation heterogeneity between samples and analyse how sampling distance, methods and sizes will impact on the ITH spatial pattern. Our results show that the sampling distance will quickly increase the ITH between samples when push rates are small, which is emphasised if the sampling size is smaller. On the contrary, high push rates will always maintain a certain level of mutation spatial mixing even with a small sampling size. These results agree with some observations in clinical data. Gates *et al*. (2019) sequenced the primary glioma biopsy samples from different distances, and compared the sample distance with the Jaccard index. Their data showed that the genetic heterogeneity increased with the sample spatial distance. To test the robustness of this observation, we applied margin-centre sampling to compare with completely random sampling. Both the ITH patterns in the centre and margin regions are consistent with our results under completely random sampling, with the centre region revealing a lower ITH compared to the margin region.

Our model provides a quantitative analysis of how growth modes, the sampling distance and size will impact on the measurements of intra-tumour heterogeneity. Those results confirm the importance of obtaining spatial information in understanding tumour evolution, as well as the possible deviation of estimated evolutionary properties such as mutation rates introduced by sampling details.

## Acknowledgements

H.L. was funded by the China Scholarship Council and Sun Yat-sen University. W.H. was funded by NSFC. B. W. was funded by the UKRI future leader fellowship. G.D. was funded by The First Affiliated Hospital of Guangzhou Medical College.

## Author contributions statement

W.H., B.W., G.D. conceived and supervised this study, H.L. W. H., and B. W. developed the methods and modelling ideas. H.Y. L. implemented computational simulations and analysis. F. T., Y. H., L.D., Z. Y., X.F. and L. W. assisted H. L. in completing parts of the algorithms. H.L., B.W. and W.H wrote the manuscript with the input of all other authors.

## Code availability

R code and Python code of full pipeline descriptions and settings used for mapping are available at https://github.com/SYSU-BioEvoLab/Spatial-Heterogeneity.

## Supplementary Information

**Supplementary Figure 1.**
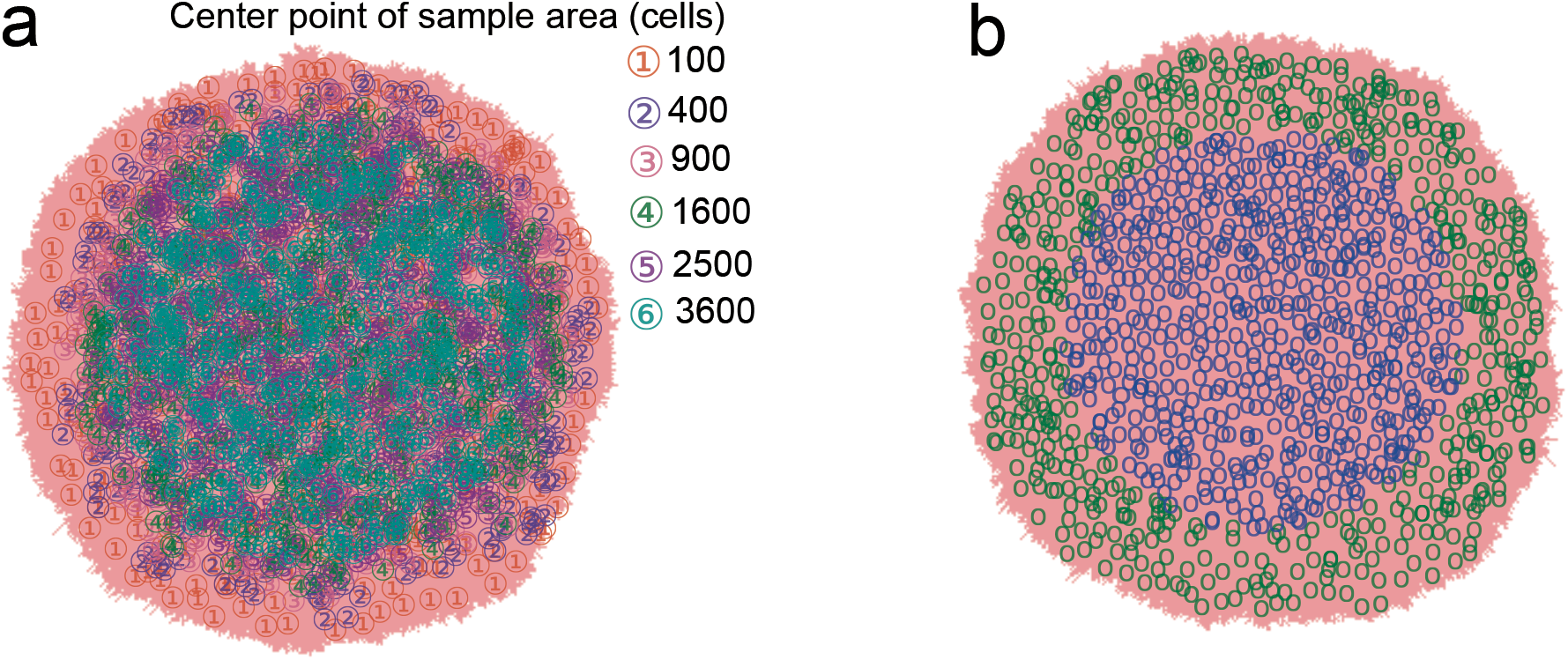
Two sampling methods, a) random sampling with different sampling sizes and b) centre and margin sampling. (1-time simulation, sampling 500 points for each size or margin and centre. tumour cell number is about 2^14^)

**Supplementary Figure 2.**
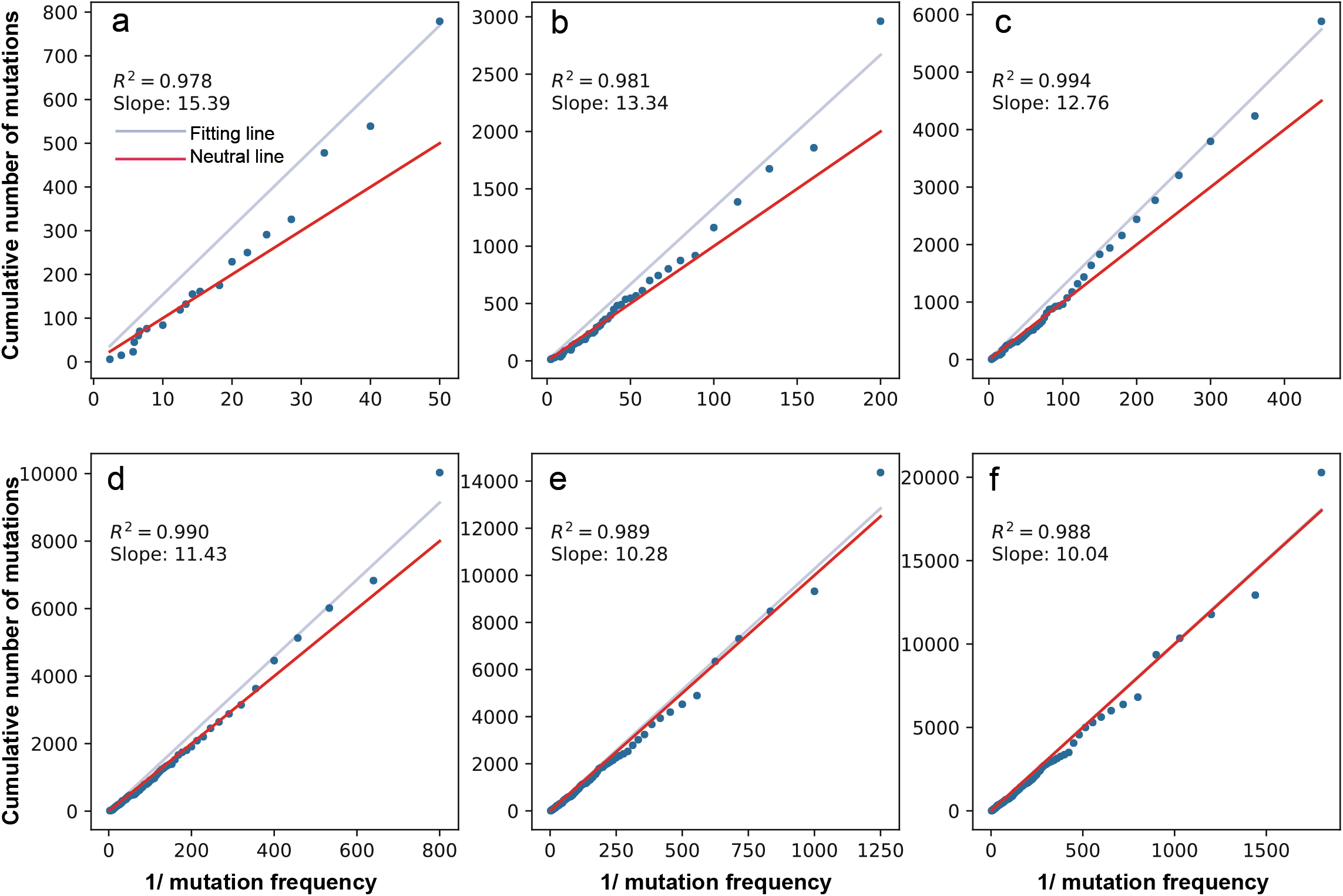
The slope and fitting level of mutation accumulation is under different sampling. (a-f: 100-3600 cells. Fit formula is *y* = *ax*. 1-time simulation, push rate=1, cell number is about 2^14^, cut last 3 points to fitting slope and *R*^2^)

**Supplementary Figure 3.**
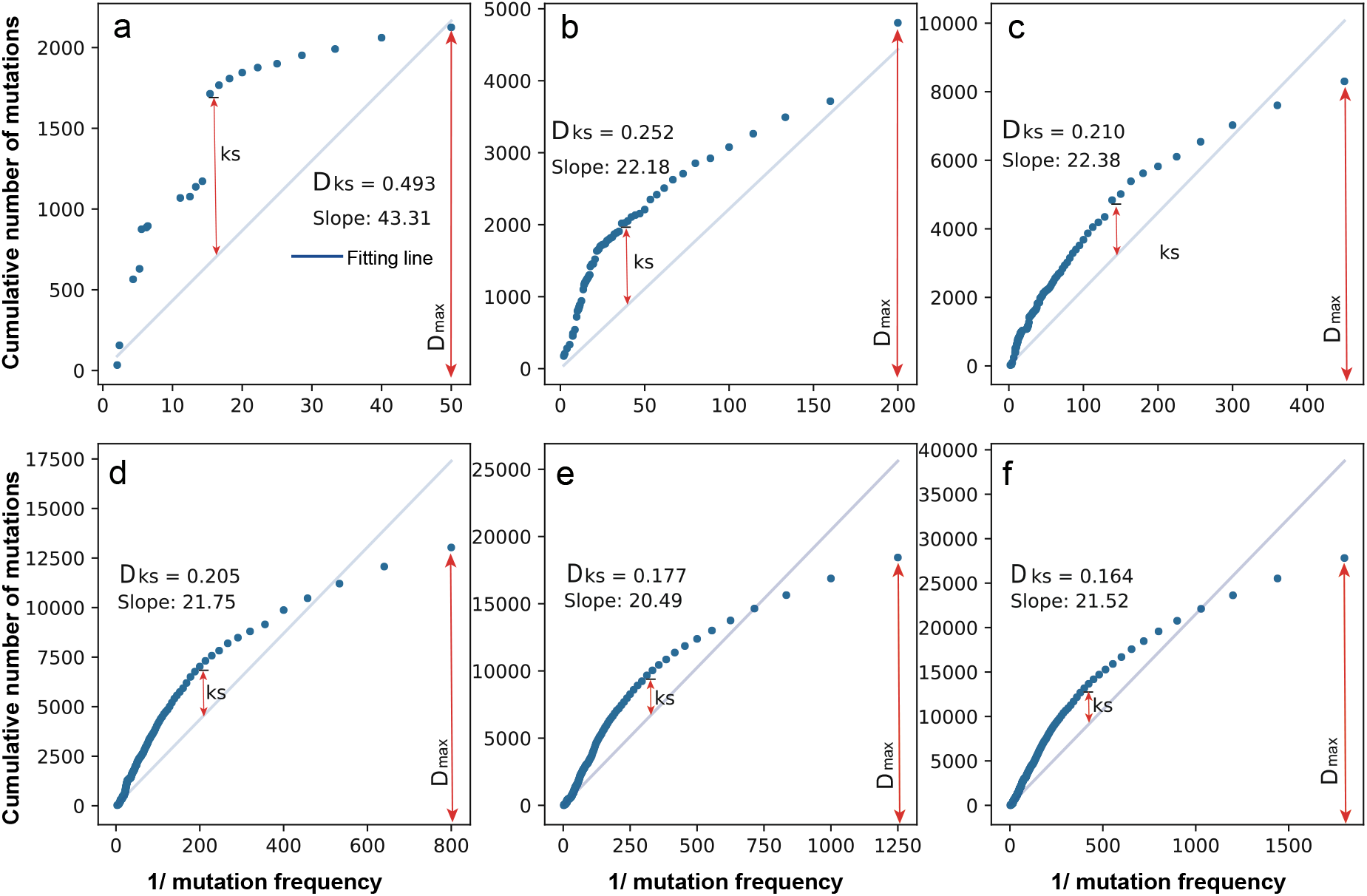
The Kolmogorov–Smirnov value between fitting line and mutation accumulation is under different sampling. (a-f: 100-3600 cells. Fit formula is *y* = *ax*. 1-time simulation, push rate=0, cell number is about 2^14^, cut last 3 points to fitting slope and *D*_*ks*_)

**Supplementary Figure 4.**
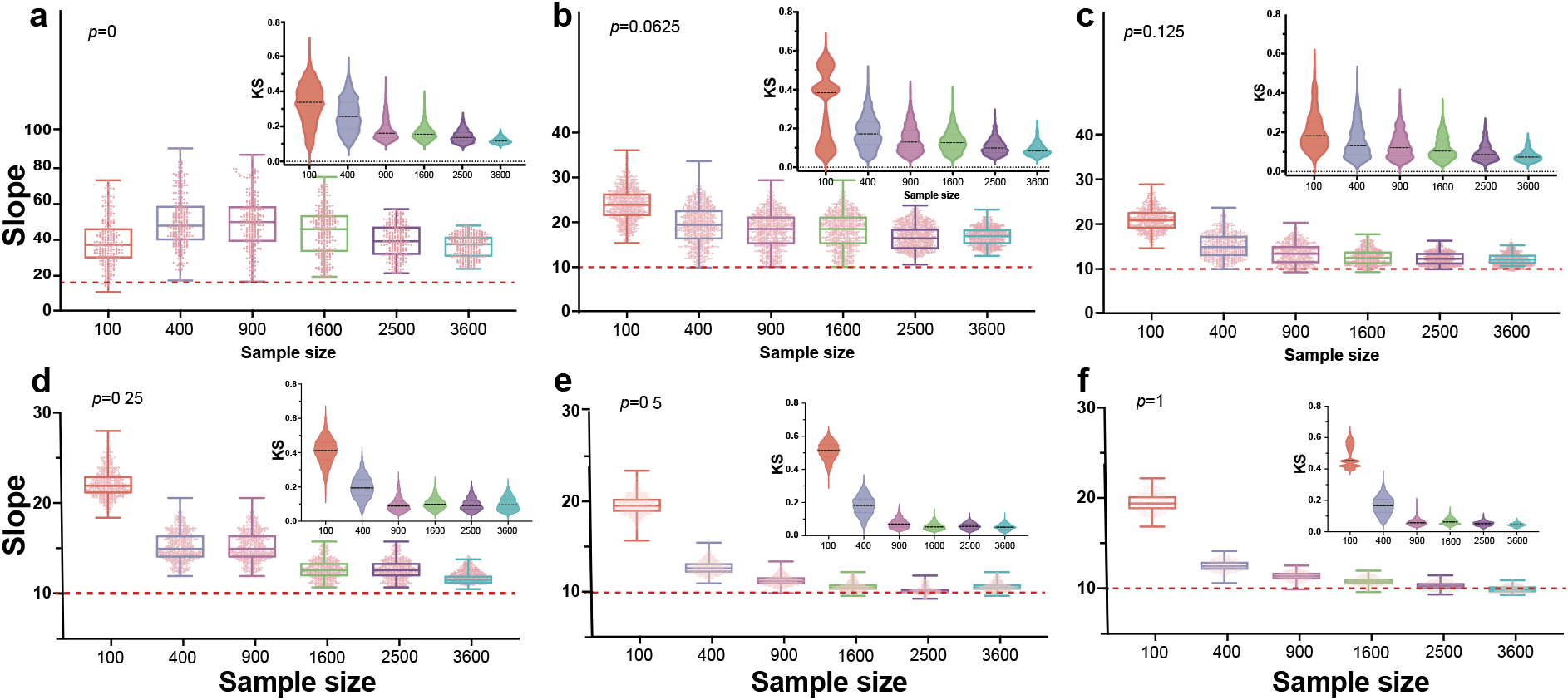
The slope fitting and Kolmogorov–Smirnov value of mutation accumulation is under different push rates. (100 times simulation, cell number is about 2^14^), cut last 3 points)

**Supplementary Figure 5.**
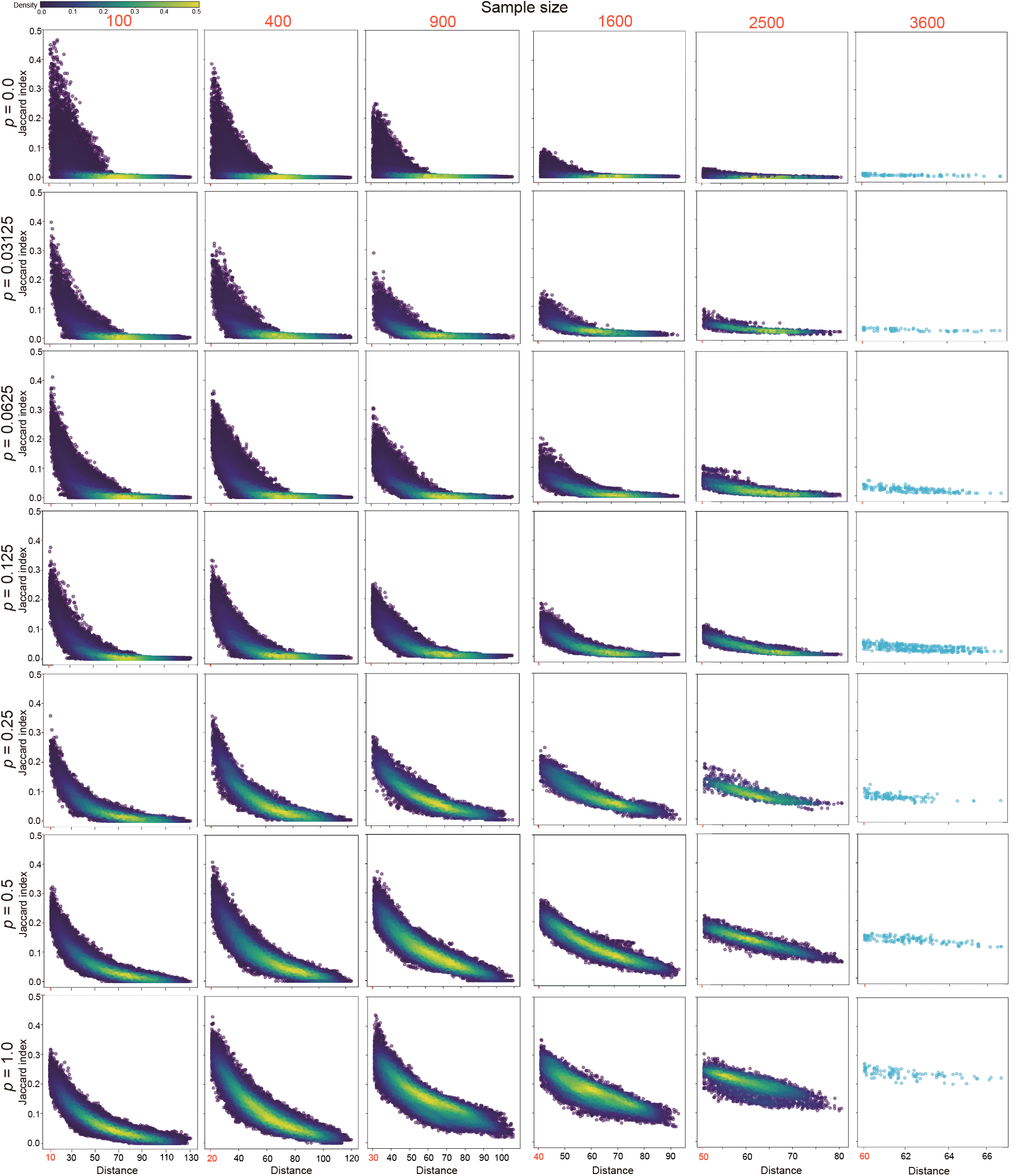
The Jaccard index distribution with non-overlapping sampling under different sampling sizes and push rates. (tumour size is 2^14^)

**Supplementary Figure 6.**
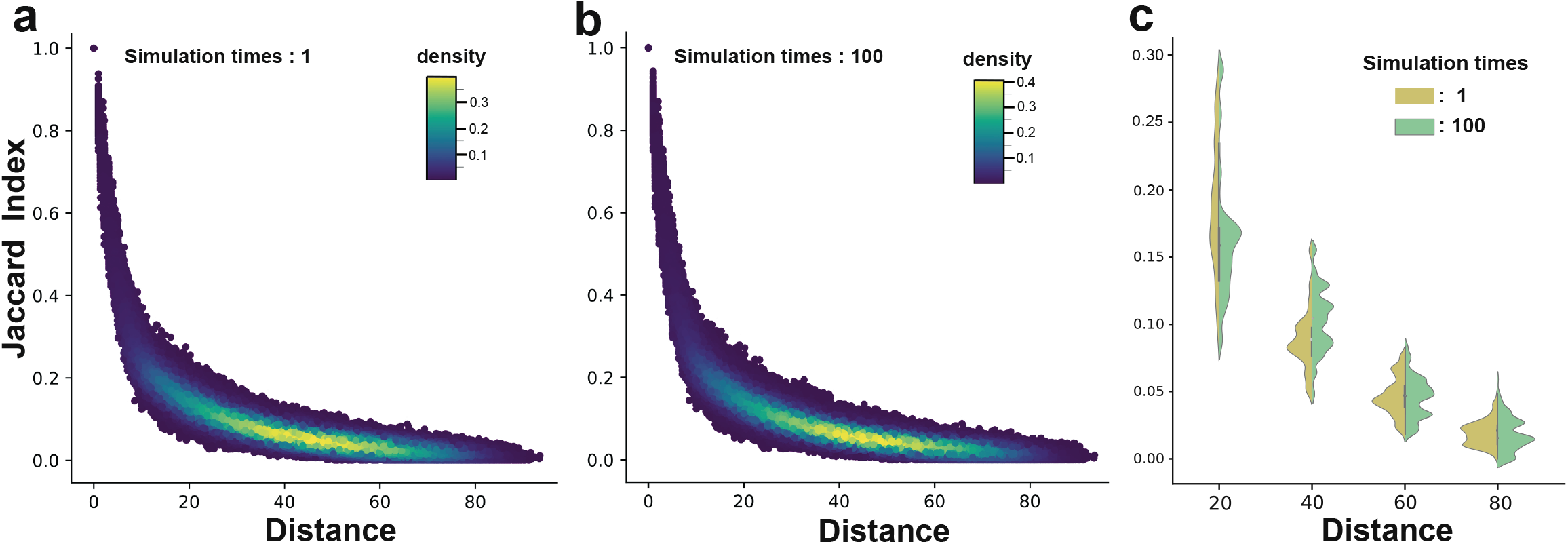
The Jaccard index distribution of 500 sampling points under one simulation and 100 simulations. (push rate is 1, the sample size is 100, tumour size is 2^14^)

**Supplementary Figure 7.**
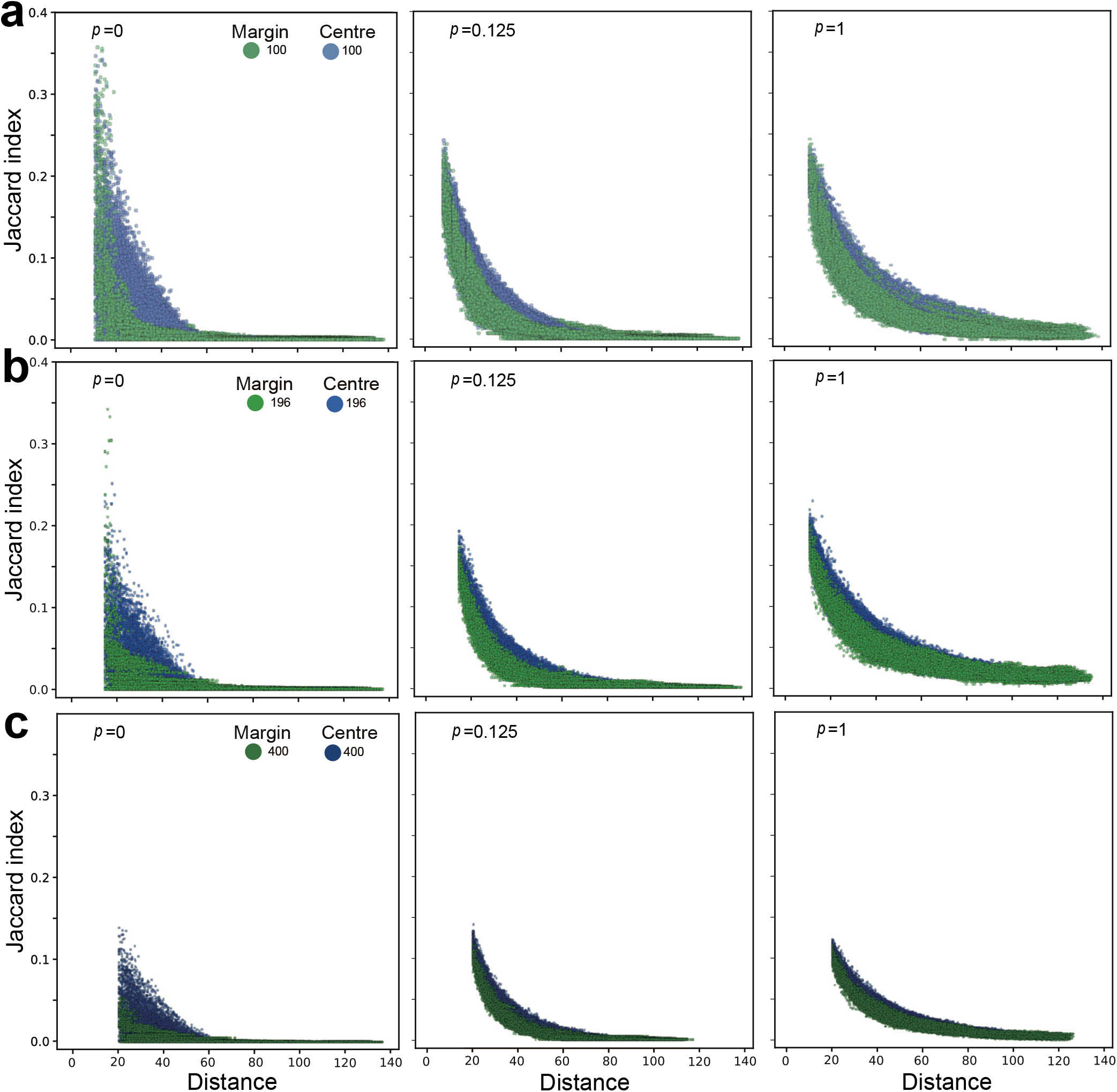
The mutation distribution of centre and margin sampling under the different push rates. **a**) with overlap points. **b**) fitter the overlap point **c**) under the different sampling sizes (1-time simulation, sampling 500 points each margin and centre, cell number is about 2^14^), cut last 3 points)

